# Genome ARTIST: a robust, high-accuracy aligner tool for mapping transposon insertions and self-insertions

**DOI:** 10.1101/024976

**Authors:** Alexandru Al. Ecovoiu, Iulian Constantin Ghionoiu, Andrei Mihai Ciuca, Attila Cristian Ratiu

**Author notes:** Corresponding author: Attila C. Ratiu.

## Abstract

A critical topic of insertional mutagenesis experiments performed on model organisms is mapping the hits of artificial transposons (ATs) at nucleotide level accuracy. Obviously, mapping errors may occur when sequencing artifacts or mutations as SNPs and small indels are present very close to the junction between a genomic sequence and a transposon inverted repeat (TIR). Another particular item of insertional mutagenesis is mapping of the transposon self-insertions and, to our best knowledge, there is no publicly available mapping tool designed to analyze such molecular events. We developed Genome ARTIST, a pairwise gapped aligner tool which works out both issues by means of an original, robust mapping strategy. Genome ARTIST is not designed to use NGS data but to analyze ATs insertions obtained in small to medium-scale mutagenesis experiments. Genome ARTIST employs a heuristic approach to find DNA sequence similarities and harnesses a multi-step implementation of a Smith-Waterman adapted algorithm to compute the mapping alignments. The experience is enhanced by easily customizable parameters and a user-friendly interface that describes the genomic landscape surrounding the insertion. Genome ARTIST deals with many genomes of bacteria and eukaryotes available in *Ensembl* and *GenBank* repositories. Our tool specifically harnesses/exploits the sequence annotation data provided by *FlyBase* for *Drosophila melanogaster* (the fruit fly), which enables mapping of insertions relative to various genomic features such as natural transposons. Genome ARTIST was tested against other alignment tools using relevant query sequences derived from the *D. melanogaster* and *Mus musculus* (mouse) genomes. Real and simulated query sequences were also comparatively inquired, revealing that Genome ARTIST is a very robust solution for mapping transposon insertions.

Genome ARTIST is a stand-alone user-friendly application, designed for high-accuracy mapping of transposon insertions and self-insertions. The tool is also useful for routine aligning assessments like detection of SNPs or checking the specificity of primers and probes. Genome ARTIST is an open source software and is available for download at www.genomeartist.ro and at www.bioinformatics.org.

## 1. Background

Consequent to the sequencing of model genomes, a massive effort was focused towards *in vivo* validation of putative genes, as an essential support for accurate biological annotations. *D. melanogaster* is arguably the most versatile eukaryote model for genetics and genomics studies and insertional mutagenesis was of paramount importance for bridging genetics and molecular genetics of this organism [Adams *et al*., 2002]. Nevertheless, many other model genomes, such as those of *Pseudomonas aeruginosa* [Jacobs *et al*., 2003], *Saccharomyces cerevisiae* [Seifert *et al*., 1986], *Caenorhabditis elegans* [Bessereau, 2006], *Danio rerio* [Kawakami, 2005] and *Arabidopsis thaliana* [Page & Grossniklaus, 2002] are also currently interrogated with transposon mutagenesis. Genetic analysis provides an impressive arsenal of tools tailored to induce, detect and characterize mutant alleles of practically any gene of interest (GOI). Although high-throughput procedures are predominant nowadays, small-scale experiments are still performed whenever particular mutant phenotypes are considered. As various alleles are required for a comprehensive functional annotation of any gene, efficient mutagens are critical tools when GOIs are considered for targeted mutagenesis. Insertional mutagenesis is a very effective strategy used to construct mutant alleles and it relies on a plethora of specific artificial transposons (ATs) designed for this purpose [Mátés *et al*., 2007; Bellen *et al*., 2011]. Many ATs are defined at their ends by TIRs, as it is the case of *P{lacW}* [Bier *et al*., 1989] and *P{EP}* [Rorth, 1996] molecular constructs, which were designed for mutagenesis of *D. melanogaster* genome. Almost all transposon insertions conduct to the duplication of a short target sequence (target site duplication or TSD), therefore each of the TIRs is flanked by a TSD [Muñoz-López & García-Pérez, 2010]. The raw data used to map the insertional mutations are query sequences containing transposon-genome junctions (or transposon-genome reads). These reads are usually obtained by sequencing specific amplicons derived by iPCR performed on DNA template extracted from specific mutants [Rehm, 2002]. Actually, mapping an insertion consists in computing the reference coordinate of the genomic nucleotide present at the juxtaposition between the genomic fragment and TIR in the transposon-genome read. We further refer to this critical nucleotide as terminal genomic nucleotide (TGN).

The mapping accuracy may be hindered when small-scale genomic mutations like SNPs or small indels are present very close to the TIR or when minor sequencing artifacts located near to the TIR affect query sequences. This issue is not manageable by available mapping tools that rely on identification and removal of the transposon fragments from the transposon-genome read. This trimming of the transposon fragments results in a shorter query sequence, which is further aligned against the reference genome, in order to identify the site of insertion. It is important to notice that, consecutive to the trimming, the impeding small-scale mutations or sequencing artifacts become located very close to the end of the new query sequence. In our mapping experience, it is a challenging issue to overpass such small-scale mutations or sequencing artifacts. Hence, the TGN is often not included in the final genomic alignment and therefore a nucleotide that precedes the mutation is erroneously reported as the insertion site instead. We developed Genome ARTIST, an application designed to map insertions of DNA entities into a reference sequence, but also the self-insertions of transposons, even when interrogated with poor-quality or mutations-bearing query sequences. The mapping strategy of Genome ARTIST is resilient to small-scale mutations and sequencing errors, providing a more accurate mapping performance as compared to similar mapping tools, such as iMapper [Kong *et al*., 2008].

Genome ARTIST is an offline, gapped heuristic aligner and it was originally conceived to map insertions of ATs in *D. melanogaster* genome using the specific files archived in FlyBase database format (ftp://ftp.flybase.net/). In order to cope with various genomes archived in Ensembl (www.ensembl.org) or NCBI (ftp://ftp.ncbi.nlm.nih.gov/) database formats, specific scripts were written in order to enable Genome ARTIST to map insertions in a wide range of prokaryote and eukaryote genomes. Herein, we describe the performances of Genome ARTIST v1.19; a less complex variant of our tool was used in previous mapping work [Ecovoiu *et al*., 2011].

## 2. Implementation

Genome ARTIST was written in C++ and JAVA for Linux OS. The minimal computer requirements are an Intel Atom 1 GHz CPU or equivalent, 1 GB of disk memory, 1 GB of RAM for bacteria and invertebrate genomes and up to 4 GB of RAM for the small vertebrate genomes. Genome ARTIST was designed for 32-bit architectures but it may also be run on a 64-bit OS version by using the detailed instructions available in Manual (available in *docs* folder). The user may either copy Genome ARTIST on the hard disk or can run it from an external device formatted as *ext3* or *ext4*. Regardless of the choice, the *Genome-ARTIST.sh* file should be selected as an executable. We tested Genome ARTIST and obtained similar performances on Ubuntu (versions 10.04, 11.04, 12.04, 13.04, 14.04), Linux Mint 14.1, Open Suse 12.3, CentOS 6.4, Fedora 19 and on Bio-Linux 8 bioinformatics workstation platform [Field *et al*., 2006]. Bio-Linux 8 is a straightforward alternative for using Genome ARTIST since it contains the pre-installed Java JDK environment and the appropriate 32-bit library required for running Genome ARTIST on the 64-bit OS version. A very convenient alternative is to avoid the installation of a Linux OS by setting two partitions on the same USB memory stick, with one bootable partition for Bio-Linux 8 and a second one containing the Genome ARTIST package. When Bio-Linux 8 runs Live from the bootable partition, the Genome ARTIST package can easily be copied on the desktop from the second partition. As a feasible alternative for the Linux environment, we tested the open-source Oracle virtual machine VirtualBox (https://www.virtualbox.org/) for emulating Bio-Linux 8 on Mac X OS and Windows platforms. Consecutive to the installation of the ISO file format of Bio-Linux 8 as a virtual machine on both OS versions, we were able to run Genome ARTIST with full performances. After opening the Genome ARTIST folder in Bio-Linux 8 environment, the user should select: *Edit > Preferences > Behavior > Ask each time* in order to customize Ubuntu 14.04 to run appropriate files as executable. The *Genome-ARTIST.sh* file must be marked as an executable following the path: *Properties > Permissions > Execute*, then Genome ARTIST can be run for mapping work. The specific scripts required to convert genome data downloaded from either Ensembl or NCBI should also be marked as executable in order to work (see Manual).

In order to compute the alignments results, different fragments of the reference sequences must be loaded in RAM, which is a time consuming step. To circumvent this aspect, the script *cachePreloadGenomes.sh* optimizes the writing of big chunks of data from the hash tables, .raw and .gene files in RAM, concomitant with launching *Genome-ARTIST.sh*. If RAM capacity is large enough to accommodate the reference genome size, the *cachePreloadGenomes.sh* completely loads the data from the respective files in the working memory, enabling Genome ARTIST to access them even easier.

## 3. The mapping strategy of Genome ARTIST

The nucleotides are binary encoded by Genome ARTIST as A = 00 (0), C = 01 (1), G = 10 (2), T = 11 (3), where the decimal conversion of binary values is shown in parentheses. Overlapped intervals of 10 nucleotides referred as decamers or *basic intervals* (*BI*) are used for indexing the reference sequences and for spanning the query sequence. During the loading of a reference AT or genome sequence, Genome ARTIST builds a hash table with an index for each decamer. The hash tables for each reference sequence are computed and saved as .*hash* files. They are accessed when interrogated with the overlapped decamers of the query sequence and then the specific addresses relative to coordinates of the reference sequences are retrieved. There are 4^10^ possible decamers in any genome and (4^10^ – 1) positions in the hash table. For example, the hash position (value) for the specific *k-mer* of 10 T nucleotides is computed by a formula considering the decimal value of T: 3×4^9^ + 3×4^8^ + 3×4^7^ + 3×4^6^ + 3×4^5^ + 3×4^4^ + 3×4^3^ + 3×4^2^ + 3×4^1^ + 3×4^0^ = 1048575, where 4^10^ = 1048576. Specific files are generated in the *resources* folder, namely distinct .*raw* files containing the standard nucleotide strand of each reference sequence and specific associated .*gene* files containing the gene annotations. By creating distinct files for each chromosome of a genome, Genome ARTIST is particularly able to work with single or many chromosomes, pending on the user choices. Genome ARTIST allows the user to customize each working session by adding or deleting chromosomes, genomes or transposons, depending on the queries or on the purposes of the research project. Obviously, the time necessary for hashing depends on the size of the genome. Multiple tests revealed that less than a minute is required for hashing a bacterial genome, a few minutes are necessary for invertebrate genomes and around 20 minutes are required for small vertebrates as *D*. *rerio* if average computing power is used. Large mammalian genomes such as those of *Mus musculus* and *Homo sapiens* are too big to be dealt with by Genome ARTIST, but either distinct chromosomes or groups of chromosomes may be loaded from any mammal reference genomes and used for mapping of insertions (about a half of the human genome is loadable in a single working package). On average, when starting a query search for a sequence of about 500 nucleotides, Genome ARTIST computes the list of the resulting alignments in a time interval ranging from seconds to tens of seconds, contingent upon the particular CPU performances and the size of the reference genome. Genome ARTIST supports mapping of multiple query sequences either in FASTA format (where care should be the taken to avoid empty spaces before the “>” symbol of the first FASTA descriptor in the list), or in text format, assuming that all query sequences in the list are separated by at least an empty row from each other.

The overlapped and/or adjacent *BIs* are merged into contiguous association intervals. Their margins are further extended by a combination of a Smith-Waterman (SW) algorithm [Smith & Waterman, 1981] implementation (SW1 step) and an original scoring formula. The expansion strategy of Genome ARTIST relies on gradually computing an alignment score for a gliding window of four nucleotides, which was designed as a robust procedure able to surpass both mutations like SNPs or small indels and various sequencing artifacts (see details in Supplementary Material 1 - SM1). The resulting product of the expansion step is referred to as an extended interval (EI) and represents an association interval between two nucleotide stretches: a query fragment and a matching nucleotide window of the reference sequence. Whenever existent, the overlapped or adjacent EIs are joined together into nucleotide associations referred as MEIs (Merged Extended Intervals). Each MEI is further converted into a proper alignment by a second SW implementation (SW2 step) and is graphically reported as a *partial alignment* (PA). Except for sequences which contain only genomic or transposon nucleotides, where the SW2 product is reported as the final result, a PA covers the query sequence just partially. All of the PAs standing for the same query sequence, regardless if they are *transposon partial alignments* (TPAs) or *genomic partial alignments* (GPAs), are reported in a single customizable list, according to the criteria of score, location or nucleotide coordinates. Each PA contains a core region referred as a *nucleus*, defined by the outermost possible lateral stretches of at least 10 consecutive nucleotide matches (see SM1). The *nucleus* is flanked by sub-alignments with lower matching density (alignment tails) and represents an item which is carefully considered during the assembly and scoring of the results. The structure and length of both the *nucleus* and the alignment tails of a PA are dependent on the settings applied for the specific parameters of Genome ARTIST (see SM1).

The main innovation of Genome ARTIST consists in the dynamic procedure used to set the border between genomic and transposon fragments present in the composite query sequences. The most challenging step of the procedure is to merge the appropriate PAs into a final alignment, in order to cover the entire query sequence and to detect the insertion coordinate with very high accuracy. To solve this item, Genome ARTIST combines TPAs and GPAs in an interactive manner, using original joining rules that govern the edge trimming and merging of PAs. The first rule is that, when overlapping, the *nucleus* of a PA is privileged over the alignment tail of the partner PA, regardless of the origin of the two PAs. A second rule is that if the *nucleus* of a TPA happens to overlap the *nucleus* of a GPA (overlapping is allowed between two *nuclei* but no more than 40% over their individual length), the shared *nucleus* fragment is allotted to the transposon in the final mapping result. This feedback between TPA and GPA entities is designed to prioritize both the TIR integrity and the structure and length of the nuclei. If the transposon fragment is not affected by mutations or by sequencing artifacts, the TIR-containing TPA would have no alignment tail towards the border with the GPA since the TPA cannot exceed the margin of the transposon reference sequence beyond the TIR. On the contrary, even when perfectly aligning composite queries are interrogated with Genome ARTIST, an alignment tail is generated at the TIR-facing end of the GPA, due to the random extension of the genomic alignment into the transposon fragment. This acquisitive behavior is possible because Genome ARTIST does not employ the standard practice of *ab initio* identification and removal of the transposon fragments to obtain cleansed genomic fragments, which are further aligned against the reference sequence. If the composite query sequence is affected by mutations or by sequencing artifacts occurring around the genome-TIR border, the alignment tails would contain them as indels and mismatches located close to each *nucleus*. It is crucial to correctly include these gaps and mismatches in the final result in order to increase the mapping accuracy. Although an intermediary TPA-GPA intersection point is estimated by Genome ARTIST, the insertion coordinate is computed only consecutive to a final re-alignment of each component PA of the final result by means of a supplemental SW adaptation. This SW3 step is applied only for those PAs which are merged into a final alignment, because the joining process often involves edge trimming of alignment tails or/and of nuclei, thus changing the context for which the alignment was optimal consecutive to SW2 step. The rationale for SW3 is simple: when mutations or sequencing artifacts are present very close to the junction border, the adjustment of the overlapped sub-alignments may affect the best possible final alignment of each modified PA, a condition which affects the mapping accuracy.

The original, key aspect of the SW3 implementation of Genome ARTIST is that the query fragment is not realigned against the exact corresponding reference nucleotide window of the PA but against a longer one. Essentially, the initial reference window is elongated with two lateral nucleotide strings, each of them representing the next 10 consecutive nucleotides of the main reference sequence. When the reference sequence window of a PA is located close to the end of the main reference sequence, one of the lateral strings is either shorter than 10 nucleotides or even absent and SW3 is accordingly performed. As a result of this approach, the gaps and mismatches located close to the border may be included in the final result. The joining strategy of Genome ARTIST overcomes mapping problems encountered when a transposon is inserted very close to SNPs or small indels in a particular genotype.

When poor quality query sequences are analyzed, false positive alignments with conjunctural better scores may obscure the actual unique insertional event. To circumvent this problem, we implemented an optional cumulative bonus score of 500, which is applicable only for alignments which contain a TIR-genome border and fulfill specific conditions described in SM1. By selectively boosting the scores of alignments that contain a TIR-genome juxtaposition, the bonus score helps the user to distinguish among real insertional events and circumstantial false positives having close aligning scores. The utility of the bonus score is evident when dealing with poor-quality query sequences which require regular/common trimming (data not shown). Genome ARTIST was devised to resolute insertions in unique genomic sequences and the bonus option is a feature supporting this purpose. On the other hand, mapping of self-insertions is a representative asset of Genome ARTIST tool and the bonus option should be avoided when mapping such molecular events. The reason is that short genomic sequences which may randomly be placed close to TIRs are highlighted if the conditions for bonus allocation are fulfilled. As a result, self-insertion of ATs (those of natural transposons as they are genomic entities) may be overlooked. Since many ATs contain in their structure genetic markers derived from the target model genome, the bonus usage may gratuitously highlight alignments which stand for apparent insertions in the corresponding genomic locations.

An example is represented by the self-insertion of *P{lacW}* construct in its own *mini-white* marker. If the bonus option is activated, the best scoring result reported by Genome ARTIST is a false positive genomic insertion in *white* locus, outscoring the real self-insertion event with the arbitrary score of 500. As a rule of thumb, whenever Genome ARTIST reports an insertion in a gene cloned in the respective AT, it is a good option to analyze the respective query sequence without the bonus option.

The mapping performances of Genome ARTIST may be fine-tuned by adjusting the values of a set of alignment parameters (detailed in SM1). Whenever illustrative for the examples described in this article, the values used to compute some particular alignments are mentioned. Technical details about the performances of Genome ARTIST are exhaustively provided in the accompanying Manual. Distinct packages of Genome ARTIST containing genomes of classical model organisms are also provided as archives at www.genomeartist.ro.

## 4. Results

The general performances of Genome ARTIST were tested with 39 original sequences derived by iPCR inquiry of *D. melanogaster* mutant strains obtained in our laboratory by mobilization of *P{lacW}* and *P{EP}* artificial transposons [Ratiu *et al*., 2008]. The trimmed sequences were deposited in GenBank database under accession numbers provided in Supplementary Material 2. These sequences represent 35 hits of *P{lacW}* and *P{EP}* in unique genomic sites, a *P{lacW}* insertion located in an *opus* transposon copy and three self-insertions of *P{lacW}.* A few of these sequences (as it is the insertion affecting *wech*) contain minor sequencing errors, a condition that makes them suitable for testing the robustness and accuracy of Genome ARTIST.

We also used Genome ARTIST to map 18 splinkerette-derived sequences from *D. melanogaster* and described in the paper of Potter and Luo [Potter & Luo, 2010]. Except for one sequence retrieved from a mutant strain having genomic features different from the reference genome, Genome ARTIST mapped these insertions in agreement with the nucleotide coordinates reported by the authors (the genome release R5.57 is used throughout this article for reporting the mapping coordinates). Additionally, we evaluated the performances of Genome ARTIST with 96 mouse-derived splinkerette sequence data made available for testing by the web page of iMapper (http://www.sanger.ac.uk/resources/software/iMapper/data/pb_test.fa). Because of the sheer size of mouse genome, we used two packages of Genome ARTIST, each loaded with about a half of the genome. All mapping results offered by Genome ARTIST were in agreement with the results computed by iMapper for these sequences. Genome ARTIST was also tested with various simulated insertional mutations affecting bacteria and various eukaryotes as *A. thaliana*, *C. elegans*, *S. cerevisiae* and *D. rerio* (data not shown); the mapping results were as theoretically expected.

### 4.1 Visualization of mapping data

Genome ARTIST offers intuitive graphical annotations such as: nucleotide coordinates for both the query and the reference sequences, the gene or the overlapped genes affected by the insertion, the left and right neighboring genes flanking the hit and the relative orientations of the transposon and genomic sequences present in the query. If present in the query sequence, the intersections of the genomic and AT fragments are presented as perpendicular borders separating blue rectangles (the genomic sequences) from red rectangles (the AT sequences). The genomic nucleotide coordinate placed immediately next to the terminal coordinate of a TIR (namely the TGN) is the critical mapping marker and Genome ARTIST reports it as the site of the insertion using blue digits. For example, the terminal coordinates of the reference sequence of *P{lacW}* construct are 1 and 10691 (FBtp0000204). Hence, the genomic reference coordinate of a TGN located consecutive either to coordinate 1 or 10961 is the one reported by Genome ARTIST as the insertion site. When any insertion occurs between two consecutive nucleotide but no TSDs are induced, two consecutive mapping coordinates may be computed, depending if the sequencing was performed at the 5’ or at the 3’ end of the insertion. On the other hand, when TSDs are generated, as it is the case for most of the described transposons [Muñoz-López & García-Pérez, 2010], an absolute mapping is not possible, as the TSD occurs both at the 5′ and the 3′ end of the insertion. Genome ARTIST do not depends on TSDs as items indispensable for mapping, even if a specific TSD may be easily inferred if both junction ends are sequenced. Although some drosophilists consider that the insertion site is represented by the first nucleotide at the 5′ end of the TSD [Bellen *et al*., 2004], any mapping convention is debatable, as correctly pointed out by Bergman [Bergman, 2012]. Actually, such an insertion is physically located between the last nucleotide of a TSD copy and the first nucleotide of the second TSD copy. Both of these nucleotides represent distinct TGNs, as each of them is proximal to a TIR. The specific TGN reported by Genome ARTIST depends on which junction end was sequenced and fed as a query sequence for aligning and mapping. The same approach is used by iMapper, which also do not consider TSDs during mapping performance. Genome ARTIST and iMapper report two different mapping coordinates when alternatively fed with query sequences standing for 5′ end and for 3′ end of the insertion. If the TSD is an octet, as it is the case for *P{lacW}*, the two coordinates are not consecutive but are separated by 7 successive positions in the genomic reference sequence. RelocaTE, a tool which uses NGS data and relies on accurate detection of both TSD copies for transposon mapping, reports two coordinates for any insertion [Robb *et al*., 2013] as, by default, there is no option to use only one end sequence/read for mapping. The two coordinates reported by RelocaTE stand for the first and respectively for the last nucleotide of the TSD, just to deal with the mapping uncertainty described above.

As an example for data visualization, we present the mapping of a *P{lacW}* insertion in *lama* gene from *D. melanogaster* (Figure 1). The blue area represents the genomic sub-sequence corresponding to *lama* while the encompassing red rectangles stand for fragments of *P{lacW}*, as in a canonical iPCR-derived sequence. The border between the terminal nucleotide of IR (coordinate 10691) and the genomic fragment reveals the site of insertion at nucleotide 5348435. The second border is at coordinate 5348475, just consecutive to GATC sequence, which represents the restriction site of *Sau3AI* restrictase used in our specific iPCR experiment, as recommended by Rehm [Rehm, 2002]. Genome ARTIST assigns the overlapped sequences to the AT, therefore *Sau3AI* restriction site sequence, which exists both in the genomic fragment and in the *P{lacW}* subsequence, is incorporated in a red rectangle.

**Figure 1.**
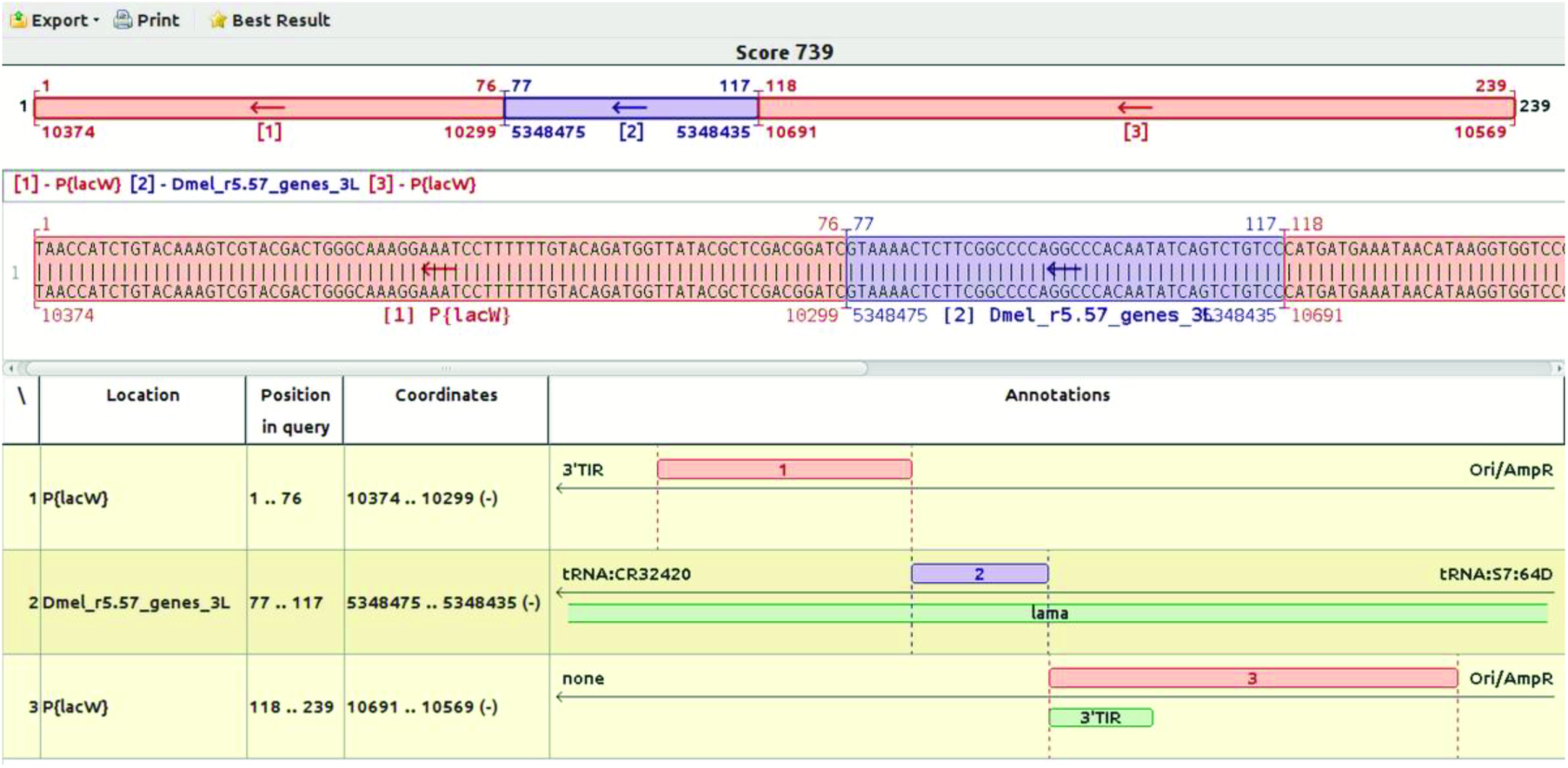
Mapping the insertion coordinate using a query sequence derived by iPCR from a *P{lacW}* hit affecting *lama* gene from *D. melanogaster*. The red rectangles stand for the transposon fragments, the blue ones represent the genomic sequence and the green ones stand for annotations of *lama* gene and of 3′ TIR of *P{lacW}*. Herein, the TGN is the C nucleotide located just next to the terminal coordinate 10691 of *P{lacW}*, which is also a C nucleotide. Hence, the insertion coordinate explicitly reported by Genome ARTIST with blue digits is 5348435. The genomic coordinate 5348475 is the one bordering the GATC restriction site of Sau3A1 used in the iPCR procedure. Since the restriction site belongs both to the transposon and to the local genomic region, it is arbitrarily allocated to the transposon sequence. Herein, we used a query sequence which contains the two transposon fragments encompassing the genomic sub-sequence.

If the genomic reference sequence files are imported in FlyBase format for *D. melanogaster*, the cytological location is shown when double-clicking on the green bar of the affected gene. Similar annotations are displayed for natural transposons or for other model genomes loaded in Genome ARTIST in Ensembl or NCBI format, excepting for the cytological coordinates.

When the coordinates of an alignment are decreasing from left to right, an arrow points to left, meaning that the graphics represent the reverse (or „-”) genomic/transposon strand and *vice versa*. There are two possible orientations of transposon insertions relative to the genomic reference strand [Bellen *et al*., 2004] and they are accordingly reported by Genome ARTIST.

The forward orientation (FWO), or orientation I, is when the reference strand of a transposon is inserted into the genomic reference strand and the reverse orientation (RVO), or orientation II, means that the reference strand of the transposon is integrated into the genomic reverse strand.

On the other hand, the sense strand of a GOI may be located either on the forward or on the reverse genomic strand and the alternatives are represented accordingly as „+” or „-”. This information is accessible in Genome ARTIST environment by double clicking on the green bar representing the affected gene. By reporting the transposon orientation relative to the sense strand of the hit gene, Genome ARTIST helps the user to check for the correct orientation of the insertion whenever a cassette-driven gene expression experiment is considered [Bellen *et al*., 2004].

Detailed instructions for interpreting the relative orientation of insertions offered by Genome ARTIST are described in Table 1.

**Table 1.**
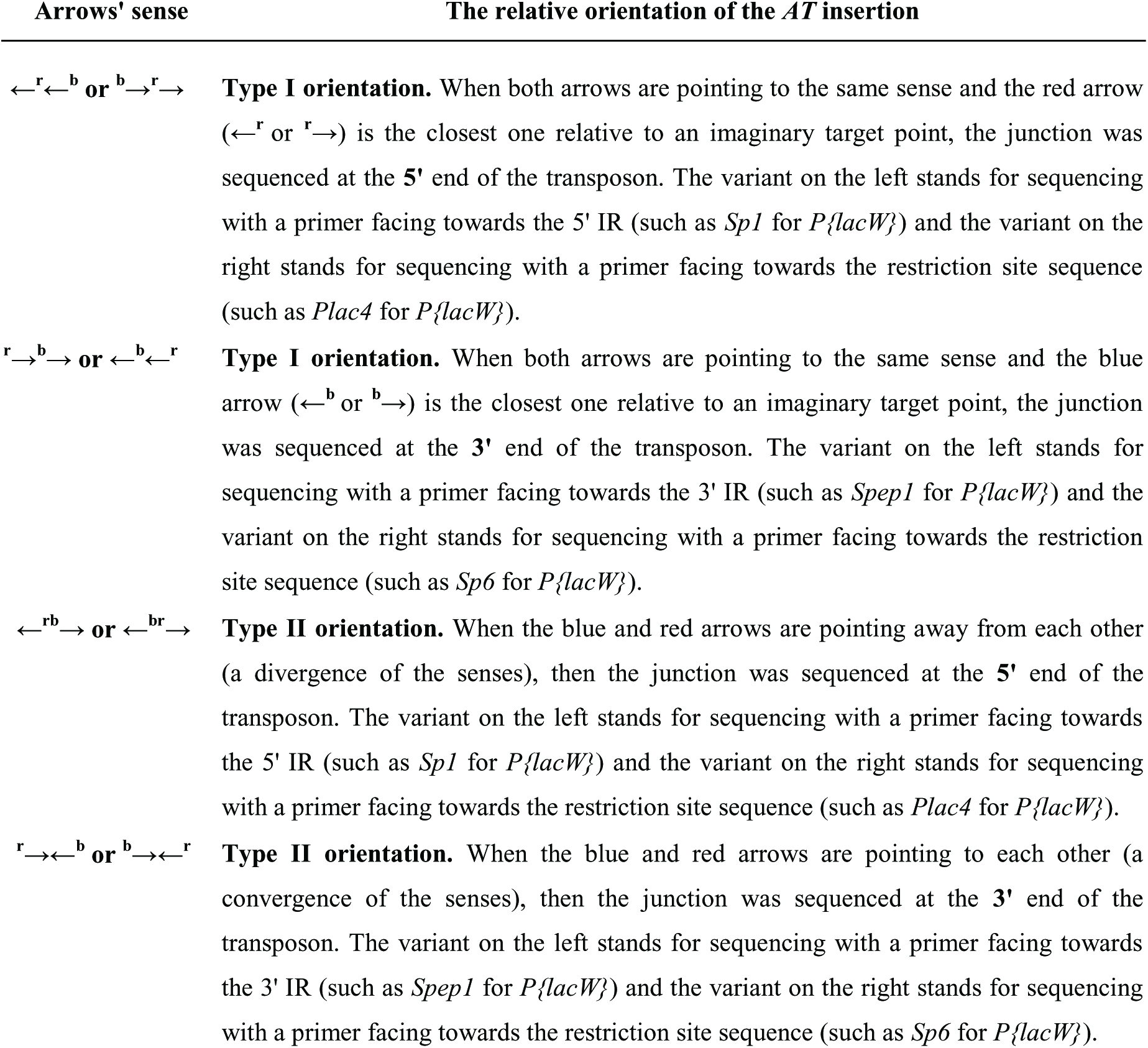
The orientation of AT insertions reported by Genome ARTIST. The superscript letter b stands for the tail of the arrow inside the blue genomic rectangle and letter r marks the tail of the arrow inside the red transposon rectangle which contains the terminal nucleotide (or the TIR). For example, the alignment for the query sequence of *P{lacW}* insertion in *lama* (Figure 1) fits the code **←^b^←^r^**. Hence, the insertion is in type I orientation and the AT-genomic junction was sequenced at the 3’ end of the transposon, with a primer facing towards the genomic sequence.

When using iMapper, only one of the two possible IRs sub-sequences may be defined as a tag, namely the one at the 3′ end of the each strand of AT, as its end points toward the genomic border of insertion. Consequently, iMapper reports as genomic sequence only the nucleotides running next to the 3′ end of the tag. Depending on the primers used for sequencing, the query sequence may need to be reverse-complemented by iMapper in order to detect the specified tag, affecting the relative orientation of the genomic sub-sequence. On the contrary, Genome ARTIST uses the entire sequence of ATs for the alignment procedure and there is no need to reverse-complements the query sequence, regardless of the primers employed for sequencing. Hence, the aligned query sequence is presented by Genome ARTIST exactly as it was entered in the search window. If necessary, a virtual iPCR sequence may be simulated by Genome ARTIST by means of a built-in option of reverse-complementing the query sequence.

Genome ARTIST displays the results as double stranded alignments, which are score-ranked in a customizable list. For each of the results, the upper strand of nucleotides represents the query sequence and the lower one contains fragments of the genomic and AT reference sequence. Due to this graphical representation, the user may also detect small mutations or polymorphisms, which are visible as mismatches or indels, a feature not offered by iMapper.

### 4.2 Mapping of self-insertions

To our knowledge, Genome ARTIST is the only available mapping tool which allows mapping of self-insertions. While other mappers trim out the AT sequences because of their potential to blur the mapping, Genome ARTIST keeps them in the query sequence. In order to compute the insertion coordinate, Genome ARTIST may use either a TIR or the whole sequence of the AT which is loaded in the transposon database. We recommend the use of the complete sequence of the AT of interest, because it allows the detection of self-insertions, aside from unique genomic insertions. Such molecular events are frequently reported for some artificial transposons [Zhang & Spradling, 1993; Golic, 1994; Spradling *et al*., 1995] and they should be accurately differentiated from genomic insertions affecting genetic markers cloned in ATs. A typical case is the one of *white* gene from *D. melanogaster*, where *mini-white* marker allele is cloned in many P element-derived constructs [Bellen *et al*., 2004]. For ATs such as *P{lacW}* and *P{EP}*, the expression of *mini-white* marker allele is essential for tracking insertional events. The graphics of Genome ARTIST enables a sharp visualization of the intersection coordinates of ATs inserted into each other. Any reference sequence, including those of ATs, may be easily annotated by the user in the Genome ARTIST environment, as it is described for *P{lacW}* in SM2. Using annotations for TIRs and genes cloned in the specific transposon permits a quick identification of the functional components affected by the self-insertion. In Figure 2, we present the case of the self-insertion event symbolized LR2.11A [GenBank: KM396322]. It may be noticed that the coordinate of this self-insertion is 8921 (as it is located just next to the terminal coordinate 1 of 5′ TIR). The self-insertion affects *mini-white* allele, therefore care should be taken not to consider it as an insertion in *white* gene located in X chromosome. Genetic analysis data revealed that LR2.11A self-insertion event is actually located on chromosome 3.

**Figure 2.**
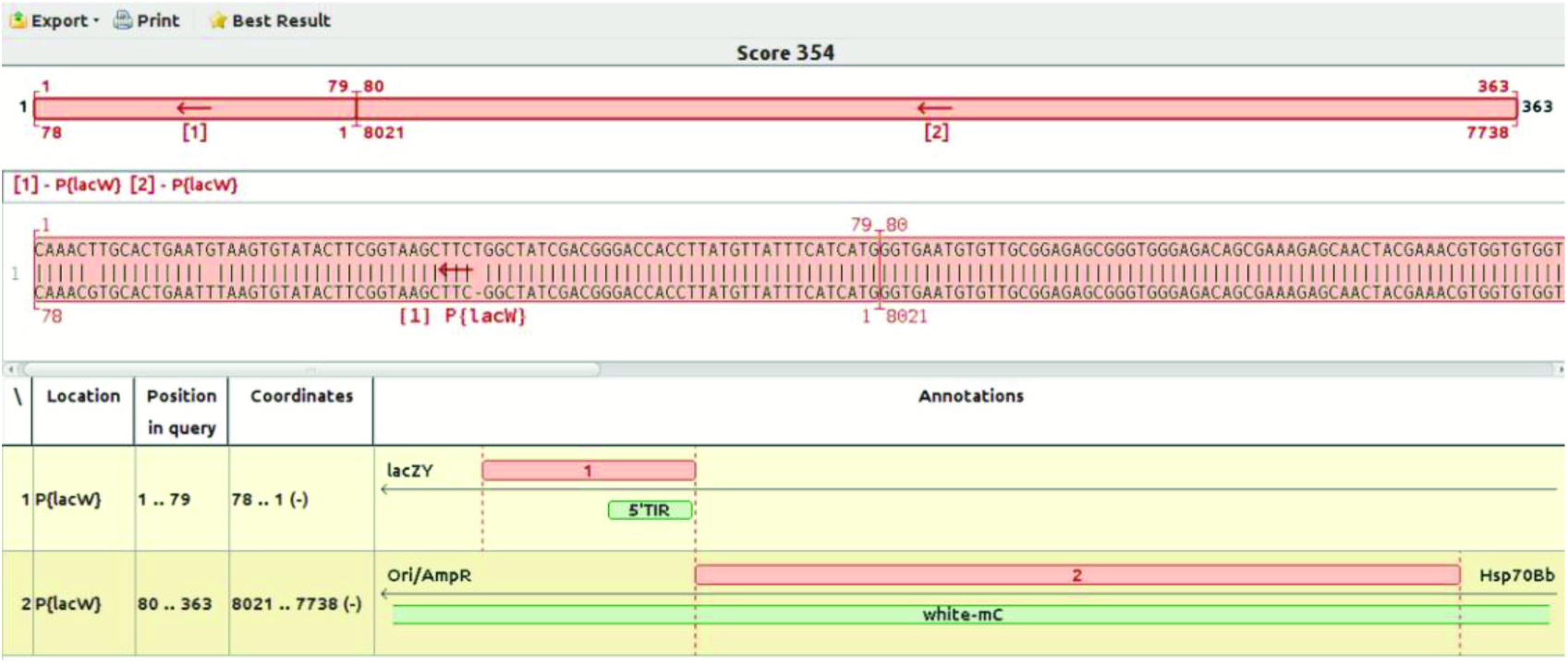
Mapping a *P{lacW}* self-insertion symbolized LR2.11A. The coordinate of self-insertion is 8921 and belongs to *mini-white* allele, which is cloned as a genetic marker in the *P{lacW}* construct.

Genome ARTIST may report marker sequences cloned in ATs as genomic fragments even when the query sequences are derived from self-insertion events. To highlight the score of a self-insertion, the bonus option should not be activated, as previously described. Mapping ambiguities specific for self-insertion events emphasize on the fact that the bioinformatics mapping data should always be correlated with the supporting genetic data.

### 4.3 Mapping insertions in particular genomic locations

According to our tests, a particular insertion of *P{EP}* construct located very close to *wech* gene *of D. melanogaster* [GenBank: GU134145] is correctly mapped by Genome ARTIST but not by iMapper, regardless the settings of its parameters. The sequence derived from the respective molecular event resolved by iPCR contains two indels in the genomic fragment as comparative to the reference sequence. As described in Figure 3, Genome ARTIST maps this insertion upstream to *wech,* at nucleotide 3377332, just next to the 3′ terminal nucleotide 7987 of *P{EP}* construct.

**Figure 3.**
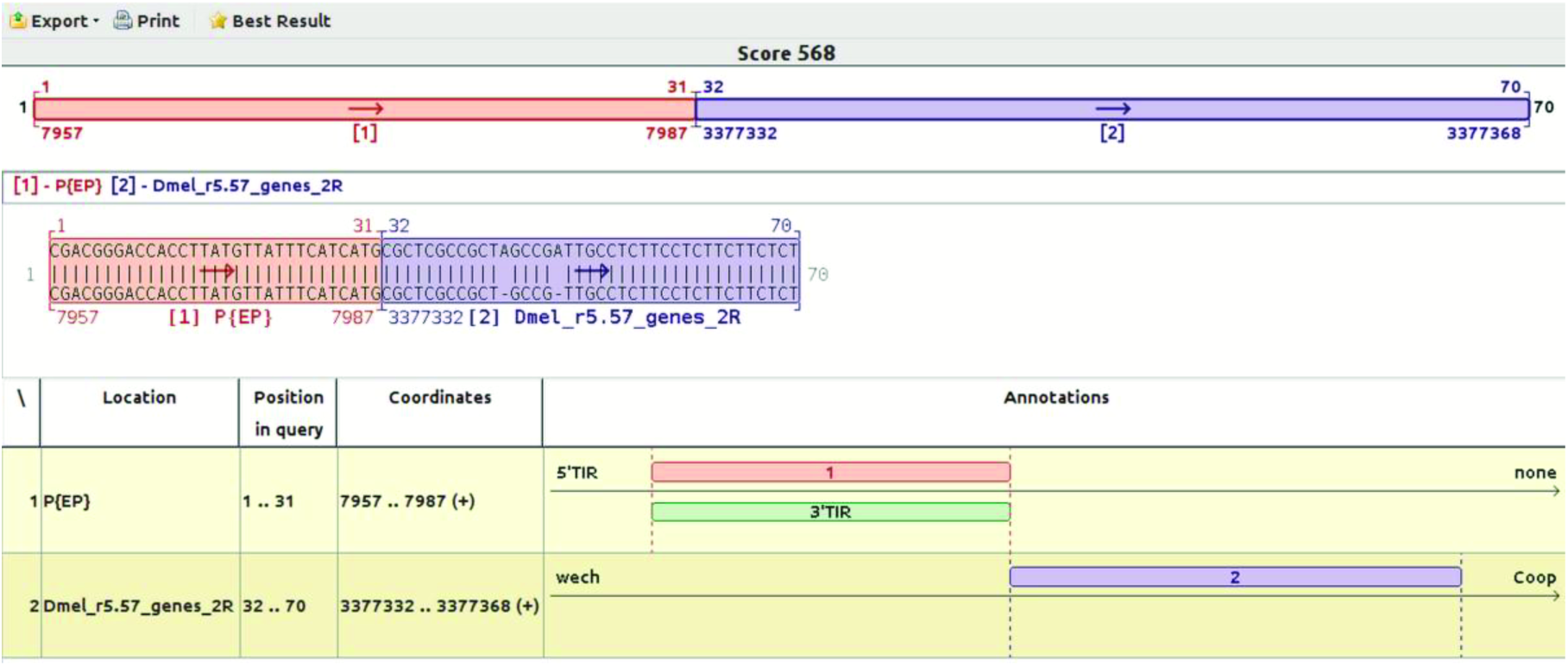
Mapping of a *P{EP}* insertion located upstream to *wech* gene. The border between the end of the *P{EP}* transposon and the genomic region points to coordinate 3377332 as the place of insertion. This coordinate is located just upstream of *wech* gene (2R) in R5.57, but in previous genome annotations it is internal to *wech* gene. The TCATG sequence present at the AT-genomic border is an overlapped sequence between the genomic fragment and the *AT* sub-sequence, but is assigned by Genome ARTIST to *P{EP}* and hence it is integrated in the red rectangle.

On the other hand, iMapper is not able to map the insertion associated with *wech,* even when the aligning parameters are set at very low stringency values. Actually, iMapper recognize the TIR as a tag, but instead reports *“No genome match found”* for the genomic sequence. The genomic fragment contains 39 nucleotides, where two supplemental adenines (As) are present as insertions relative to the reference sequence. We trimmed the sequence in order to eliminate the insertions, but iMapper is still unable to recognize the genomic sequence of 37 consecutive matching nucleotides. When the genomic sub-sequence was artificially elongated from 37 to exactly 57 nucleotides of reference *wech* sequence (and the two inserted adenines are trimmed out), iMapper was able to report the correct coordinate of insertion upstream of *wech*. If the two adenines are kept, *wech* sequence has to be elongated from 39 to 83 nucleotides, regardless of the parameters’ settings.

It is interesting to interrogate why iMapper does not recognize the string of 37 consecutive matching nucleotides upstream of *wech*. Most probably, this situation reflects a lower sensitivity of SSAHA aligner as comparative to the aligning heuristic of Genome ARTIST. As described by the authors [Ning *et al*., 2001], SSAHA constructs the hash table by searching only for non-overlapped *k-tuples* (equivalent to *words* or *k-mers*), whereas Genome ARTIST considers overlapped *k-mers* for the hash table. Additionally, SSAHA excludes from the hash table the words having a frequency above a cutoff threshold N, in order to get rid of hits matching repetitive sequences. It may be noticed that the genomic sequence of *wech* query sequence contains a CT-rich fragment (Figure 3), therefore SSAHA implementation used by iMapper may consider this sequence as containing a repetitive pattern. The example of *wech* insertions points the fact that insertions in specific regions of the reference genome may be lost if a mapper is not designed to detect even problematic insertions. The laboratory practice evidences that iPCR technology often generates short genomic sequences depending on the position in the reference genome of a specific restriction site relative to the TIRs; the closer the restriction site, the shorter the genomic fragment in the *iPCR* amplicon.

Whenever an IR terminal sub-sequence incidentally overlaps a genomic sub-sequence in a specific query, the superimposed fragment is reported as pertaining to the genome by either online BLAST [Altschul *et al*., 1990] or BLAT [Kent, 2002], since the reference sequences of ATs sequences are not compiled in the reference genomes. Therefore, the user may erroneously infer that the insertion site is located next to the overlapped fragment if the result is not manually annotated. As an example, the critical sub-sequence TCATG present in query sequence derived from the *wech* mutant is an overlap between the terminal nucleotides of *P{EP}* and the genomic nucleotides interval 3377327-3377332. If *P{EP}* construct is present in the database of Genome ARTIST, our application interprets the overlapped sequence as belonging to the IR of *P{EP*} and accurately reports 3377332 as the site of insertion. On the contrary, BLAST and BLAT algorithms erroneously report the coordinate 3377327 as the insertion point. Even more confusing, the best alignment scores reported by either online BLAST or BLAT for this query do not refer to *wech* but to paralogous HSP genes (3R).

### 4.4 Mathematical simulation of sensitivity – Genome ARTIST versus BLAT as aligners

As an aligner, Genome ARTIST resembles BLAT regarding to hashing and searching strategy, but there are relevant differences between the two aligners. For example, BLAT builds an index of non-overlapping 11-mers of the genome [Kent, 2002], while Genome ARTIST generates an index for all of the possible overlapped 10-mers that may occur in a genome. This indexing strategy enables Genome ARTIST to compare each overlapped 10-mer of the query with each overlapped 10-mer of the reference genome, which results in a higher sensitivity of aligning (here we define sensitivity as the ability of finding the maximum number of possible alignments in a given homologous interval). To compare the sensitivities of Genome ARTIST and BLAT, we employed a mathematical simulation relying on tabular data published for BLAT [Kent, 2002] and an adaptation of a mathematical recursive formula which may be found at http://www.askamathematician.com/2010/07/q-whats-the-chance-of-getting-a-run-of-k-successes-in-n-bernoulli-trials-why-use-approximations-when-the-exact-answer-is-known/ address on “Ask a Mathematician/Ask a Physicist” site. For specific homologous intervals having the same range of percentage identities (p.i.) and sequence sizes (N) we compared both algorithms and the results are presented in Table 2 and Table 3.

**Table 2.** The sensitivity of Genome ARTIST algorithm calculated when various p.i. values of homologous intervals (defined as percentage identities) of size N (ranging from 10 to 100 nucleotides) are considered. The length of the perfect match is 10 bp and the number of overlapping 10-mers in the homologous regions is N – 10 + 1.

| p.i. | 10 | 20 | 30 | 40 | 50 | 60 | 70 | 80 | 90 | 100 |
| --- | --- | --- | --- | --- | --- | --- | --- | --- | --- | --- |
| **81%** | 0.122 | 0.353 | 0.531 | 0.661 | 0.755 | 0.823 | 0.872 | 0.908 | 0.933 | 0.952 |
| **83%** | 0.155 | 0.419 | 0.610 | 0.739 | 0.826 | 0.884 | 0.922 | 0.948 | 0.965 | 0.977 |
| **85%** | 0.197 | 0.492 | 0.690 | 0.812 | 0.886 | 0.930 | 0.958 | 0.974 | 0.984 | 0.991 |
| **87%** | 0.248 | 0.571 | 0.767 | 0.874 | 0.932 | 0.963 | 0.980 | 0.989 | 0.994 | 0.997 |
| **89%** | 0.312 | 0.655 | 0.838 | 0.925 | 0.965 | 0.984 | 0.992 | 0.997 | 0.998 | 0.999 |
| **91%** | 0.389 | 0.740 | 0.899 | 0.961 | 0.985 | 0.994 | 0.998 | 0.999 | 1.000 | 1.000 |
| **93%** | 0.484 | 0.823 | 0.946 | 0.984 | 0.995 | 0.999 | 1.000 | 1.000 | 1.000 | 1.000 |
| **95%** | 0.599 | 0.898 | 0.978 | 0.995 | 0.999 | 1.000 | 1.000 | 1.000 | 1.000 | 1.000 |
| **97%** | 0.737 | 0.959 | 0.995 | 0.999 | 1.000 | 1.000 | 1.000 | 1.000 | 1.000 | 1.000 |

**Table 3.**
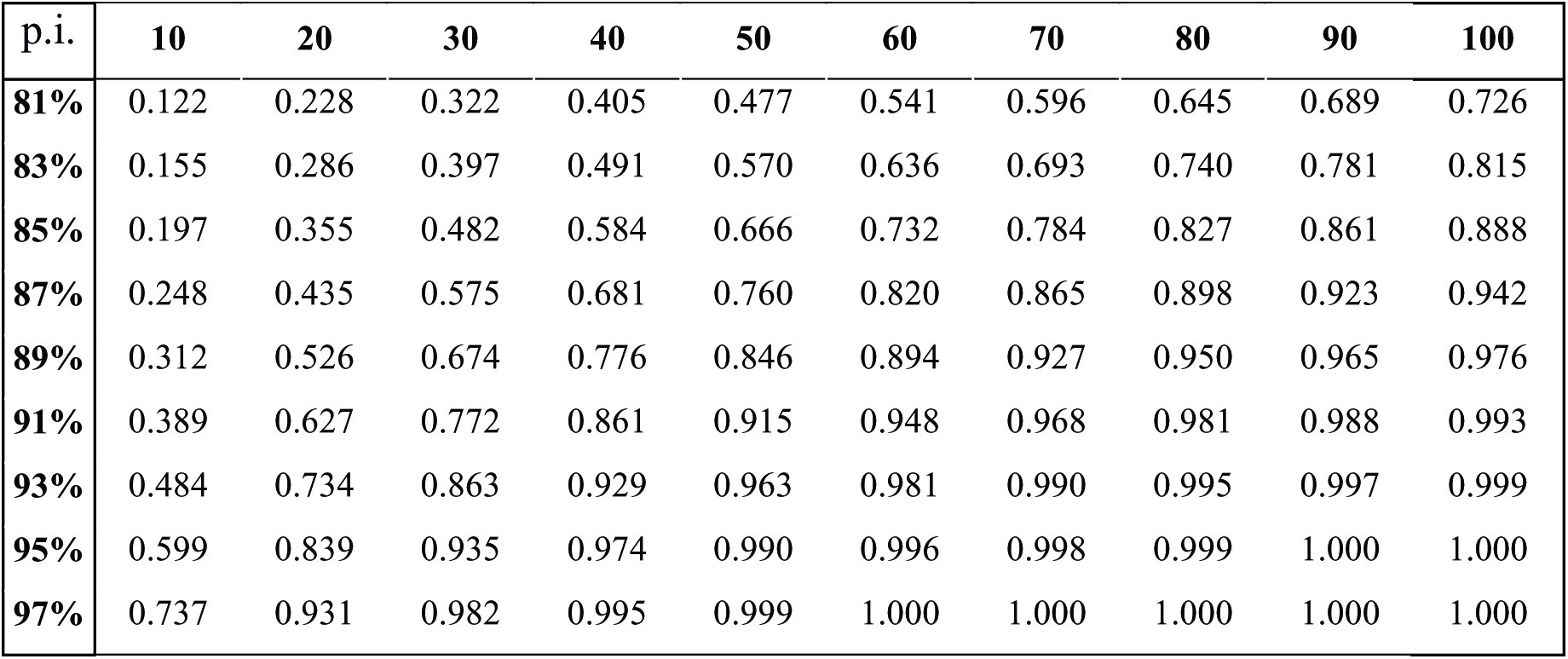
The sensitivity of Genome ARTIST algorithm when various p.i. values of homologous intervals (defined as percentage identities) of size N (ranging from 10 to 100 nucleotides) are considered. The length of the perfect match is 10 bp and the number of non-overlapping 10-mers in the homologous regions is [floor (N/10)].

A graphical representation of the values corresponding to N = 50 indicates that the sensitivity of Genome ARTIST is higher when the homology is lower than 93% (Figure 4A). Similar, for an arbitrary value of p.i. = 87%, Genome ARTIST is more sensitive than BLAT when query sequences of 20-100 nucleotides are searched against the database (Figure 4B), the most evident differences being calculated when considering 30 to 50 nucleotide long intervals. As query sequences shorter than one hundred of nucleotides are often obtained by sequencing of iPCR products derived from insertional mutants, it appears that Genome ARTIST is better suited for the alignment of such queries.

**Figure 4.**
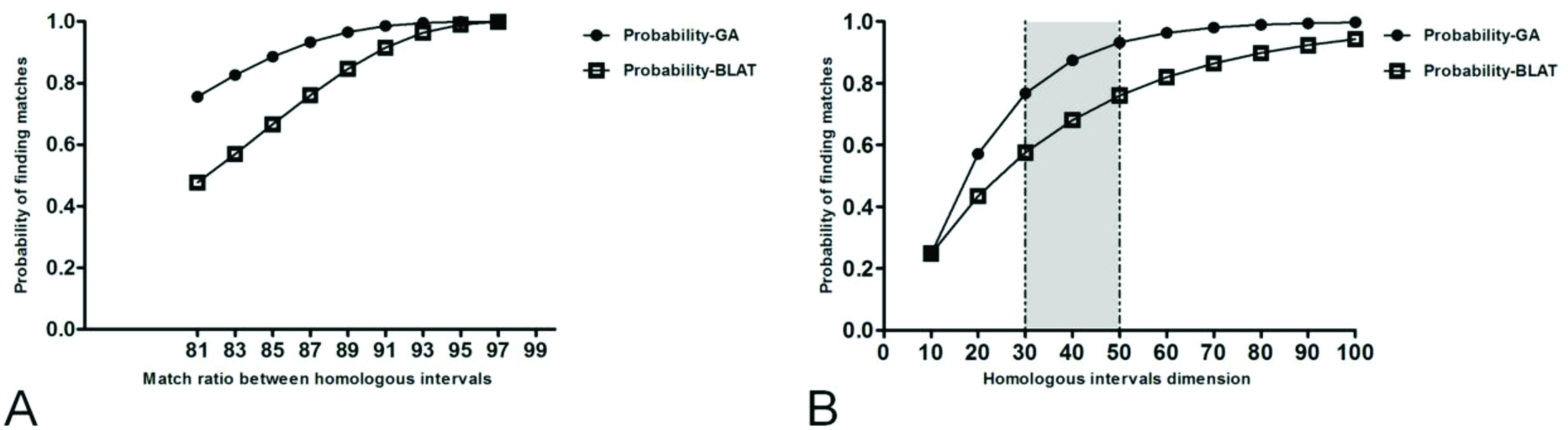
The sensitivity of Genome ARTIST compared to the sensitivity of BLAT. Figure 4A stands for a graphical representation of the probability to find in a 50 bp homologous target interval at least one overlapped (the case of Genome ARTIST) or one non-overlapped (the case of BLAT) 10-mer which perfectly matches a 10-mer from the query, when the match ratio between homologous intervals ranges from 81% to 99%. The results, represented in circles for Genome ARTIST and in squares for BLAT, reveal that the overlapped 10-mers approach enables Genome ARTIST to be more sensitive than BLAT as the match ratio decreases towards 80%. Figure 4B shows the probability to find at least one overlapping or non-overlapping 10-mer which perfectly matches a 10-mer from the query in homologous intervals ranging from 10 to 100 bp. An arbitrary match ratio of 87% between the query and each of the target homologous intervals was considered for exemplification. When the size of the homologous intervals ranges from 30 bp to 50 bp, a maximum difference in the probability of finding perfect matches is observed (gray area). The higher sensitivity of Genome ARTIST is a consequence of employing overlapping decamers during the alignment procedure (figures were generated using GraphPad Prism version 5.04 for Windows, GraphPad Software, La Jolla California USA, www.graphpad.com).

### 4.5 Genome ARTIST is robust to small-scale mutations and sequencing artifacts

When small-scale mutations (polymorphisms) or sequencing artifacts reside close to TIR-genome junction, the robustness and accuracy of the mapping tool is essential for the precise mapping of the insertion. Herein, we comparatively test Genome ARTIST versus iMapper when feeding both tools with the same query sequences. We used 23 sequences derived by iPCR from real insertions of *P{lacW}* in *D. melanogaster* genome (the original sequences are available in NCBI/EBI/DDBJ under accession numbers indicated in Supplementary Material 2). Genome ARTIST successfully mapped all the insertions with *Short* option and the bonus 500 assigned (the recommended parameters), while iMapper with default parameters is able to map 22/23 insertions to the same coordinates mapped by Genome ARTIST. The exception stands for *CR43650* gene sequence [GenBank: HM210947.1], where the value of iMapper parameter *SSAHA mapping score* should be slightly lowered from >35 to >34 in order to obtain a correct coordinate of insertion.

To test the mapping robustness of both Genome ARTIST and iMapper tools to small-scale mutations or sequencing errors, we handled all of the 23 sequences in order to place SNPs and small deletions inside a presumptive TSD of 8 nucleotides. The range of mutated interval starts with the second nucleotide closest to the TIR and ends at the 6^th^ nucleotide outside of the TIR, as described in Figure 5.

**Figure 5.**
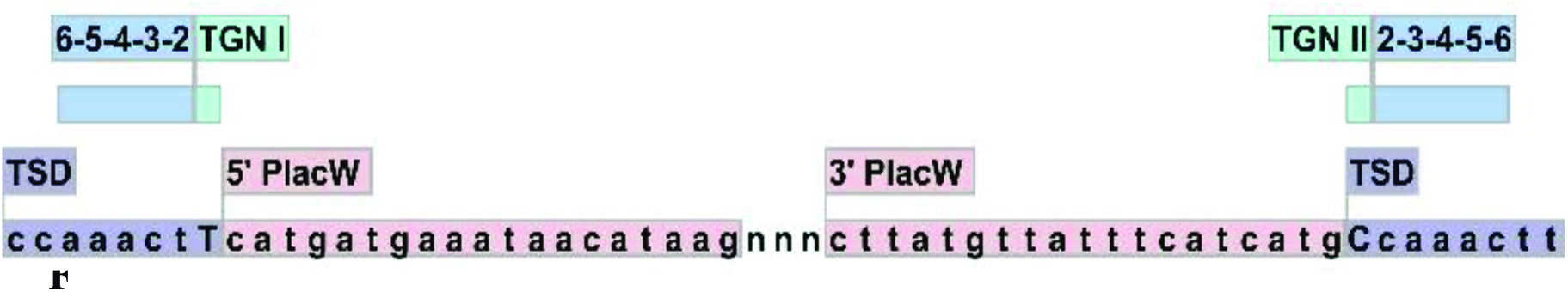
Simulation of small-scale mutations affecting nucleotides located close to the TIR, in a region equivalent to TSD, which is represented herein by the arbitrary octet CCAAACTT (blue). With reddish are highlighted the partial sequences of the two TIRs specific for *P{lacW}* construct. TGN I (a T nucleotide) and TGN II (a C nucleotide) are capitalized inside the respective TSD boxes. The nucleotides affected by simulations in TSD are those located in the relative positions 2, 3, 4, 5 and 6 as sliding away from each TGN toward the other end of TSD. The drawing was realized with CLC Main Workbench software v.6.9 (CLC Bio-Qiagen, Aarhus, Denmark).

The patterns of simulations for each of the 23 sequences were generated in a step by step approach. As a result, we induced SNPs affecting positions 2, 3, 4, 5 or 6 relative to TGN; one-nucleotide deletions affecting positions 2, 3, 4, 5 or 6 relative to TGN; substitutions of two consecutive nucleotides simultaneously affecting positions 3 and 4 relative to TGN; deletions of two consecutive nucleotides simultaneously affecting positions 3 and 4 relative to TGN; substitutions of three consecutive nucleotides simultaneously affecting positions 3, 4 and 5 relative to TGN; deletions of three nucleotides simultaneously affecting positions 3, 4 and 5 relative to TGN. We always kept the TGN unmodified since it should be reported as the genomic coordinate of the insertion if the simulated small-scale mutations are properly overpassed.

We noticed that, when affected, the most sensitive positions of TSD are 2, 3 and 4, as they impede the mapping accuracy of both Genome ARTIST and iMapper. Nevertheless, Genome ARTIST still reports the real insertion coordinates for most of the sensitive simulations, reflecting the ability of our tool to surpass small-scale mutations occurring very close to the TIR. In our hands, iMapper fails to report the real coordinate of insertions for all sets of 23 simulations which affect nucleotide 2, 3, and 4 (except for the SNP affecting nucleotide 4), even when the mapping parameters were set for the most permissive values. The comparative results of mapping the simulated sequences are presented in Table 4 and in Figure 6.

**Table 4.** The accuracy of the mapping results reported by Genome ARTIST and iMapper for various patterns of simulated small-scale deletions and SNPs in sets of 23 real query sequences.

| Position of simulated mutation in TSD | Deletions (precise mappings) |  | SNPs (precise mappings) |  |
| --- | --- | --- | --- | --- |
|  | Genome ARTIST | iMapper | Genome ARTIST | iMapper |
| 2 | 14 | 0 | 16 | 0 |
| 3 | 20 | 0 | 23 | 0 |
| 4 | 23 | 0 | 21 | 21 |
| 5 | 23 | 3 | 23 | 21 |
| 6 | 23 | 20 | 23 | 22 |
| 3 and 4 | 17 | 0 | 17 | 0 |
| 3, 4 and 5 | 14 | 0 | 12 | 0 |

**Figure 6.**
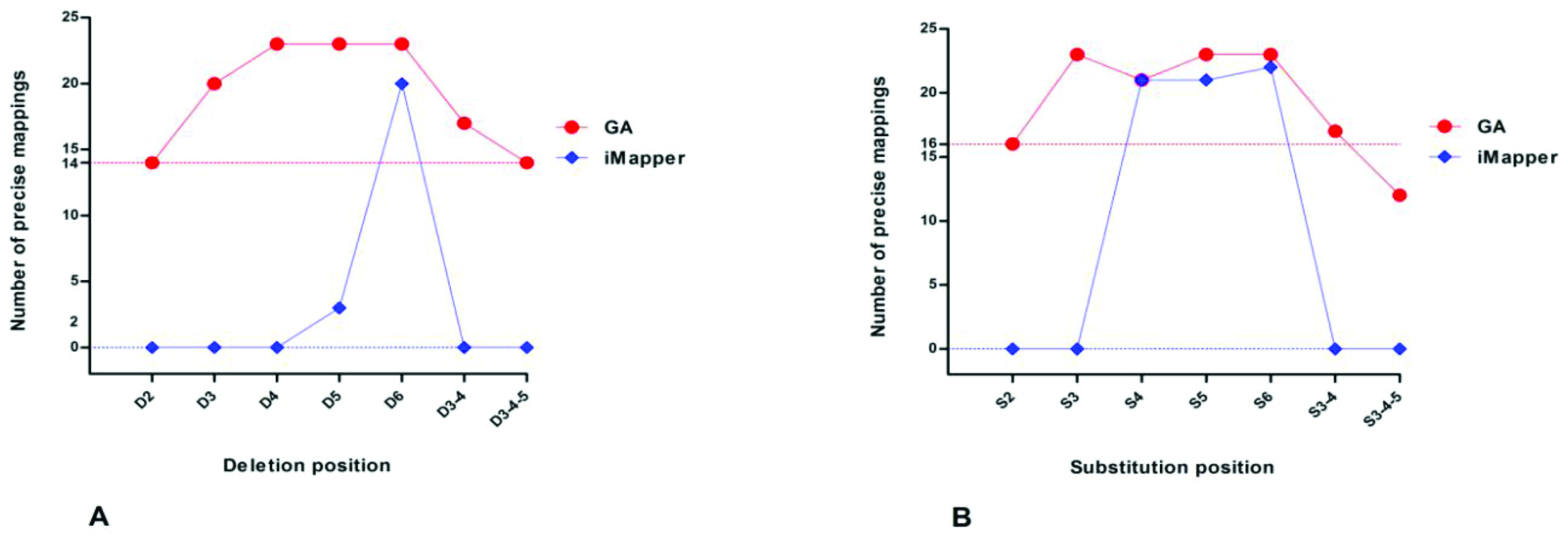
A comparison of Genome ARTIST (in red) and iMapper (in blue) mapping performances when small-scale mutations are simulated as either deletions or substitutions in a set of 23 original query sequences obtained by insertions of *P{lacW}* in the genome of *D. melanogaster*. Genome ARTIST correctly maps 14/23 insertions (y-axis) when one-nucleotide deletion in position 2 relative to TGN is simulated (x-axis) and 16/23 insertions if a SNP is placed in the same position (the most critical one for mapping accuracy), while iMapper is unable to accurately map any of the respective simulated insertions. For the majority of the other simulations, Genome ARTIST is also superior to performances of iMapper.

A real issue is the fact that, although it highlights the alignment details for the TIR fragment of the query, iMapper does not present the pair-wise alignment of the genomic fragment, which actually contains the TGN standing for the coordinate of insertion. Actually, iMapper graphically displays the genomic sub-sequence of the query in a rather mechanistic manner. As a result, whenever mutations occur close to TIR-genome junction, the insertion coordinate reported by iMapper may not be the one corresponding to the nucleotide depicted as bordering the junction. In other words, as presented in Figure 7, the intuitive TGN is not the same with the nucleotide standing for the site of insertion. On the contrary, Genome ARTIST offers explicit graphics of each sub-alignment and unambiguously displays the TGN, an approach which is useful when polymorphisms or sequencing artifacts are present in the query sequence. The coordinate of insertion reported by Genome ARTIST is always the same with the graphically visible TGN.

**Figure 7.**
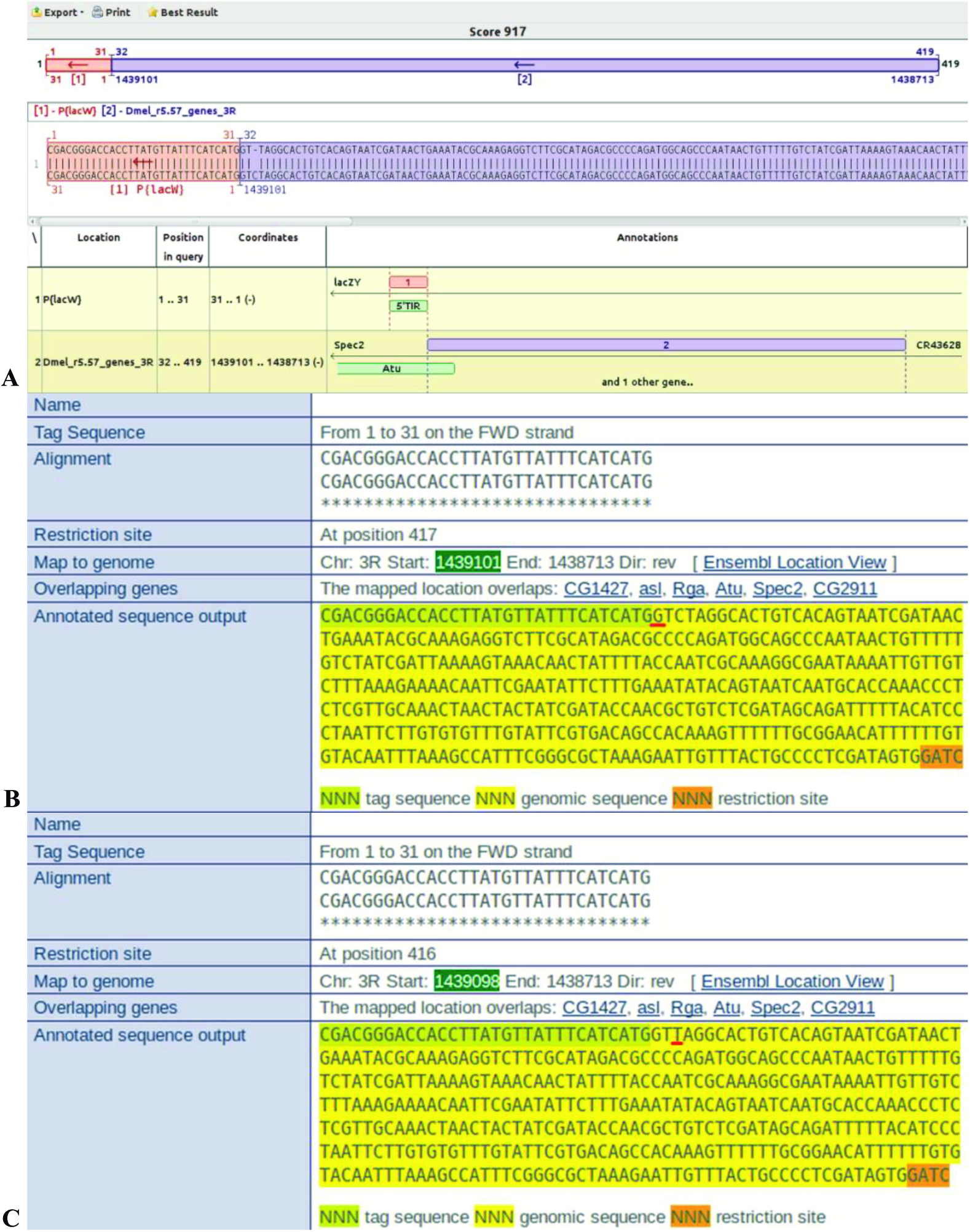
A deletion affecting the genomic nucleotide 3 relative to the TIR was simulated for *a P{flacW}* insertion affecting *Atu* gene from *D. melanogaster* [GenBank: HM210951.1]. While Genome ARTIST still reports the coordinate 1439101 as the insertion site (Figure 7A), Mapper erroneously displays the T (underlined in red) at the coordinate 1439098 (highlighted with green) as the mapping coordinate for the simulated query sequence (Figure 7C). Nevertheless, the respective T nucleotide is not intuitively positioned right next to the terminal nucleotide of the TIR (or the tag sequence, highlighted with light green by Mapper). Instead, a G nucleotide, which actually should stand for the real mapping position at 1439101 (Figure 7B), is somehow confusingly depicted as the one bordering the TIR. Moreover, the deleted nucleotide is explicitly represented by Genome ARTIST, but it cannot be directly inferred from the graphics of Mapper.

Our results reveal that Genome ARTIST is more tolerant than iMapper to small-scale mutations and sequencing artifacts residing near the transposon-genome junction. The analysis of our simulations pointed that the first three nucleotide of the TDS located just next to the TGN (as described in Figure 5) are critical positions for the mapping accuracy. When mutagenized, these positions are interpreted by Genome ARTIST rather as a buffer zone, favoring a robust detection of the TGN’s coordinate. Genome ARTIST is able to accurately deal with both small-scale mutations and sequencing artifacts, mainly due to its expansion procedure and to the interactive strategy of joining TPAs and GPAs. Obviously, the complex procedure that enables the accurate joining of transposon and genomic fragments would not be possible if the transposon fragments are removed from the composite query. Actually, this commonly employed approach would reduce Genome ARTIST to a mere aligner tool. The sheer attempt of Genome ARTIST to cover the entire composite query sequence by a best-scoring final alignment is a premise for the TPA-GPA merging step. This joining operation triggers the SW3 step, which reconsiders some nucleotides removed by edge trimming of TPAs and GPAs, but which are actually crucial for the mapping accuracy. As a result of SW3 step, some key nucleotides placed around the T-G border, including the TGN, are ultimately incorporated or rearranged in the final alignment if the TSD or the TIR are affected by mutations or sequencing errors. Genome ARTIST also applies SW3 step for other less common, but possible junctions, such as TPA-TPA and GPA-GPA ones.

We also analyzed the 23 simulated sequences containing SNPs and one-nucleotide deletions affecting the critical positions 2 and 3 of the TSD, using as an *ad hoc* mapper a BLAST implementation executed by CLC Main Workbench 6.9 software (CLC Bio-Qiagen, Aarhus, Denmark). Both stringent (word = 10; match = 1, mismatch = -4, gap opening cost = 5, gap extension = 2) and permissive (word = 4; match = 1, mismatch = -1, gap opening cost = 0, gap extension = 2) settings were used. BLAST accurately mapped a maximum of 11 simulated sequences (less than 50% of the tested sequences), regardless of the aligning parameters’ settings (data not shown). As expected, BLAST performed less precise when the stringent settings were used. For the stringent settings, only 1/23 precise mapping coordinates were obtained for the deletion affecting position 2, but 0/23 correct results were obtained when the deletion affected position 3 relative to the TIR. When the permissive settings were used for the same deletions, 2/23 correct results were obtained for both variants of simulations. Regarding to SNP simulations, 0/23 proper results were obtained for both SNP positions with stringent settings, but when using permissive settings 0/23 and 11/23 real mapping coordinates were obtained when the SNP affected nucleotides from positions 2 and 3, respectively. These data reveal a higher accuracy of Genome ARTIST when mapping insertions starting from simulated query sequences, as compared to the BLAST version implemented by CLC Main Workbench 6.9 software.

The alignment extension specific to Genome ARTIST allows the correct detection of the TGN in many of the simulated sequences even when the TIR was trimmed out. In our hands, such a performance was not attainable with either BLAST or BLAT aligners when considering the same simulations (data not shown). It appears that SSAHA, BLAST and BLAT aligners are facing problems when dealing with insertion sequences containing terminal small-scale mutations whenever the transposon sequences are removed from the composite query. Therefore, we consider that Genome ARTIST is a particularly robust alternative as both an aligner and a mapper for problematic query sequences.

## 5. Discussions

To test the mapping performances of various tools, the simulations of transposon insertions in the target genome is a current practice [Zhuang *et al*., 2014]. Instead of using this kind of experimental modeling, we simulated genomic small-scale mutations very close to the TIRs of 23 real *P{lacW}* insertions located in *D. melanogaster* genome. This approach was intended to comparatively test the robustness of Genome ARTIST to map ATs insertions when affected by polymorphisms and/or by sequencing artifacts as compared to the similar achievements of iMapper, BLAST and BLAT. According to our results, the accuracy of insertion mapping is affected when mutations or sequencing artifacts are present around the TIR-genome border or when repetitive patterns occur in the genome fragment of the query sequence. Genome ARTIST is able to surpass these problems, as revealed by the simulations of small-scale mutations data and by the *wech* example. Therefore, the robustness of Genome ARTIST represents a real advantage when such query sequences are inquired for mapping of insertions. Apart from the set of simulated sequences totaling 23 × 7 = 161 variants (Table 4), we also comparatively mapped a number of 153 unhandled insertions and Genome ARTIST always detected the right insertion coordinate.

Self-insertions are molecular events reported for artificial transposons in classical studies [Spradling *et al*., 1995]. To our knowledge, Genome ARTIST is the only tool able to map both self-insertions and genomic insertions of ATs, but mapping of natural transposons is also feasible. As the natural transposons represent a very consistent fraction of the EK genomes [Feschotte & Pritham, 2007] an application able to annotate insertions relative to both targeted genes and to natural transposons is of practical interest for this research field. In Figure 8, we present relative mapping data of a real *P{lacW}* insertion in a copy of *opus*, a natural transposon from *D. melanogaster* [GenBank: KM593302.2]. Which copy of *opus* is actually affected may eventually be revealed only consecutive to applying a PCR splinkerette procedure to the mutant line.

**Figure 8.**
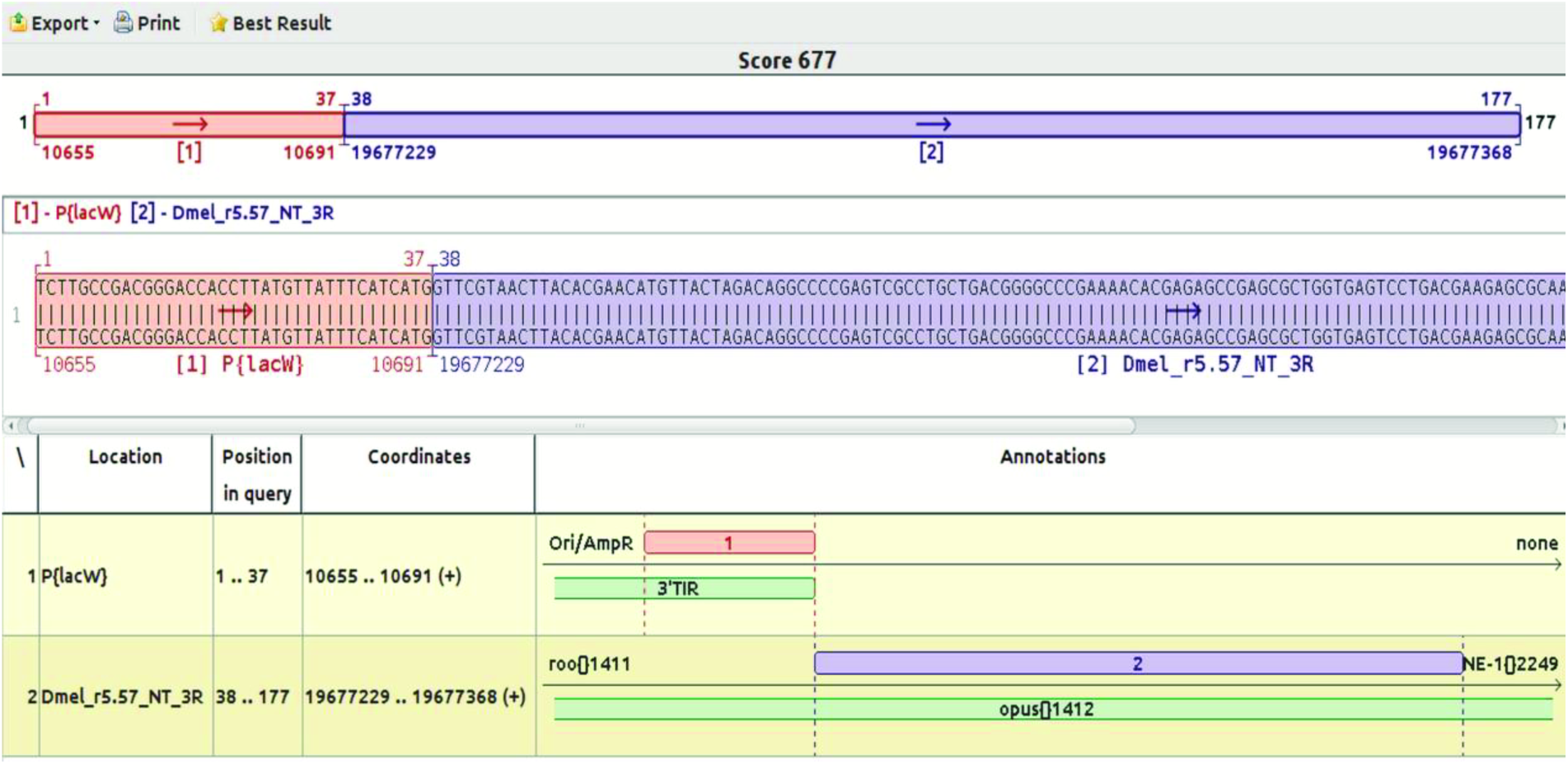
A *P{lacW}* hits a copy of the natural transposon *opus[]1421* from *D. melanogaster*. This relative result may be visualized if Genome ARTIST employs the specific annotations for natural transposons, which are available on FlyBase (see Manual). The coordinate 19677229 stands for a possible site of insertion, as many copies of *opus* are present in the genome of *D. melanogaster*. When using a query sequence derived by *splinkerette* PCR, Genome ARTIST should provide mapping coordinates for a unique, specific *opus* copy.

Multimers of transposons may be generated by nested transpositions or by self-insertions when copies of a transposon hit the original insertion of the respective mobile element [Jiang & Wessler, 2001]. This insertional behavior is a driving force for genome evolution as described in maize [Jiang & Wessler, 2001] and *D. melanogaster* studies [Bergman *et al*., 2006]. Therefore, mapping of self-insertions is of particular interest for experiments aiming to decipher the biological significance of nested transposition phenomena.

As an online application, iMapper works only with a few predefined animal genomes from *Ensembl* repository (an exception is the *S. cerevisiae* genome). Supplemental genomes may be added upon request, according to the authors [Kong *et al*., 2008], but only from *Ensembl* repository, which may be a limiting option. As a difference, Genome ARTIST deals with a broader spectrum of genomes, ranging from those of bacteria to the ones of small vertebrates. The only prerequisite is the availability in the public databases of the annotated sequenced genomes in formats that may be converted with the accompanying scripts of Genome ARTIST (see Manual). Additionally, Genome ARTIST allows the user to load and annotate genomic and/or transposon reference sequences, as described in Supplementary Material 3 and in the Manual. We successfully tested Genome ARTIST with the genomes of *P. aeruginosa*, *S. cerevisiae, C. elegans*, *D. rerio* and *A. thaliana*.

An additional advantage of Genome ARTIST is the fact that different releases of a genome may be co-loaded in the same package to test for inherent differences of annotations. The user of Genome ARTIST may work either with a whole genome of interest or with individual chromosomes, since the conversion scripts generate the output in such a way that individual chromosome files may be selected (see Manual). If short orthologies are to be hunted, small and medium size genomes of different species may be simultaneously interrogated with the same query sequence. Similarly, if various ATs are employed in an insertional mutagenesis experiment, all of their reference sequences may be co-loaded in the Genome ARTIST database.

RelocaTE [Robb *et al*., 2013], ngs_te_mapper [Linheiro & Bergman, 2012] and TIF [Nakagome *et al*., 2014] tools were designed to employ TSDs to map transposons when starting from split-reads (junction reads) obtained by NGS sequencing. A split-read or a junction read contains a fragment of the inquired transposon linked to a unique genomic fragment. The TDSs are detected and then used for merging unique genomic subsequences into small contigs which are further aligned with either BLAST (TIF) or BLAT aligner (RelocaTE and ngs_te_mapper) to find the mapping coordinates. Both TIF and RelocaTE report both terminal coordinates of the detected TSD as the insertion site, as revealed in a comparative work of mapping insertions of *Tos17* transposon in *ttm2* and *ttm5* lines of *japonica* rice cv. Nipponbare [Nakagome *et al*., 2014].

Mapping of transposon insertions consecutive to targeted PCR and Sanger sequencing versus mapping when starting from NGS data are different endeavors, a reality reflected in the algorithms developed to cope with this mapping strategies. The split reads obtained by NGS are short and more prone to sequencing artifacts, hence both high sequencing coverage and detection of perfectly overlapping TSDs are ideally required for mapping insertions at nucleotide level accuracy. On the contrary, the junction sequences obtained by the robust Sanger method starting from amplicons generated by inverse PCR or by vectorette PCR are more reliable. These sequences are, on average, an order of magnitude longer (hundreds of nucleotides instead of a few tens as in NGS). They contain unique genomic fragments embraced by two molecular markers, namely a TIR and the restriction site used for cutting the genomic DNA of the insertional mutant. In these cases, sequencing of genomic sequences flanking both ends of the inserted AT (which, indeed, would allow to confirm the TSD presence) is recommended, but not mandatory for an accurate mapping. In our experience, the detection of the two TSD copies is not a critical aspect *per se* when mapping insertions starting from PCR amplicons as it is when using short split-read sequences obtained in NGS projects. Moreover, it is notorious that sometimes sequencing at both ends of the insertion is quite difficult because of technical reasons [Ashburner *et al*., 2005; Ecovoiu *et al*., 2009]. Hence, sequencing a genomic region flanking only one end of the AT should be enough as long as either the derived sequence is of high quality or the bioinformatics mapping tool used to interpret it is very accurate. Genome ARTIST is not depending on TSDs detection for mapping and successfully deals with query sequences affected by sequencing artifacts or with small polymorphisms occurring very close to the TIRs.

Tangram uses split-reads obtained by NGS for precise mapping of insertions and implements SCISSORS program to find the breakpoint between the transposon sequence and the genomic one [Wu *et al*., 2014]. As a drawback, the authors mention that mapping errors may occur when transposon and genomic sequences are similar. According to the authors, Tangram’s analysis may conduct to erroneous mapping results when short sequences from split-reads are common to both genomic and transposon sub-sequences. The algorithm used by Genome ARTIST for computing the precise border between transposon and genomic sub-sequences of a junction sequence circumvents this problem by always assigning the overlapped sequences to TPAs and, implicitly, to the TIR. This strategy is designed to cover the whole junction query sequence by a single, final alignment, an original approach which provides very accurate mapping performances.

According to our tests, Genome ARTIST may also be used to map insertion sites of integrative viruses, as herpes simplex virus. Such a task can be easily accomplished if the virus reference sequence is loaded into the transposon database of Genome ARTIST. Depending on the genes affected by the virus integration, accurate mapping could be of biological or medical relevance. Additionally, Genome ARTIST offers very reliable results when used for SNP detection or when checking the specificity of oligonucleotides (as primers and probes) against a reference genome. The field of transposon mapping software is heavily relying on Linux environment as revealed by the fact that some recent transposon mapping tools are actually developed for Unix/Linux. Relevant examples are represented by software/programs like TEMP [Zhuang *et al*., 2014], TIF [Nakagome *et al*., 2014] and ITIS [Jiang *et al*., 2015]. Genome ARTIST is an open-source software which runs on many flavors of Linux OS and perfectly fits the popular BioLinux8 workbench.

## 6. Conclusion

Genome ARTIST is a very robust and accurate software designed for mapping insertions and self-insertions of ATs occurring in transposon mutagenesis experiments. BLAST (implemented by CLC Main Workbench v6.9), BLAT (implemented by RelocaTE and TEMP), SSAHA (implemented by iMapper), BWA (ngs_te_mapper) and Bowtie (implemented by TAPDANCE) [Sarver *et al*., 2012] are very efficient pairwise aligners, but none of them was specifically designed for mapping transposon insertions. Particularly, Genome ARTIST mapper consists in an original pairwise aligner and a particular algorithm designed to accurately join the TPAs and GPAs. This mapping strategy provides a high tolerance to small-scale mutations and sequencing artifacts occurring at the junction region between transposon and genomic sub-sequences as compared to the similar iMapper tool. Genome ARTIST is a very tweakable tool and is not dependent on permanent Internet connection, as long as a genome-loaded package of Genome ARTIST is equivalent to a backup of the respective genome data.

## Availability and requirements

Project Name: Genome ARTIST (ARtificial Transposon Insertion Site Tracker)

Project Home Page: the source code of Genome ARTIST is accessible at www.bioinformatics.org and various working packages are available for download at www.genomeartist.ro.

Operating System: Linux OS

Other requirements: JAVA JRE and one of *lib32z1 lib32ncurses5 lib32bz2-1.0* or *libstdc++6:i386*

License: GNU General Public License

Restrictions of use by non-academics: None

## Authors’ contributions

AAE conceived the concept of Genome ARTIST and coordinated the project. ICG and AMC wrote the source code. ICG wrote the conversion scripts for Ensembl and NCBI genome files. AAE and ACR extensively tested Genome ARTIST for mapping performances. AAE searched Genome ARTIST for bugs. AAE and ACR equally performed the mapping work with unhandled and simulated query sequences to comparatively test the mapping robustness of Genome ARTIST. AAE tested the software on various distributions of Linux OS. The manuscript was written by AAE and ACR and edited by ICG and AMC. AAE and ICG wrote the Manual. AMC manages the web page www.genomeartist.ro

## Acknowledgements

This work was partially supported by CNCSIS research project 147/2007, PNII-IDEI Program, granted to A.A.E, Department of Genetics, Faculty of Biology, University of Bucharest, Romania.

